# Machine Learning-Driven Identification of Virulence Determinants in *Borrelia burgdorferi* Associated with Human Dissemination

**DOI:** 10.1101/2025.07.09.663762

**Authors:** Hoa T. Nguyen, Catherine A. Brissette

## Abstract

Lyme disease, the most common tick-born infectious diseases in the United States, presents with highly variable clinical outcomes, ranging from localized erythema migrans to severe disseminated complications affecting the heart, joints, and nervous system. The bacterial determinants underlying this phenotypic variation remain largely unknown, limiting our ability to predict disease progression and optimize treatment strategies. Here, we applied machine learning (ML) approaches to identify specific amino acid residues within surface-exposed virulence factors that predict human dissemination phenotypes. Utilizing the whole genome sequences from 299 clinical Bb isolates, we extracted and characterized variants of seven known virulence factors (BB_0406, BBK32, DbpA, OspA, OspC, P66, and RevA). Protein variants were classified based on their association with disseminated versus localized infections using clinical metadata. Cramér’s V analysis revealed strong associations between dissemination phenotypes and five adhesins: BBK32, DbpA, OspC, P66, and RevA. We developed ML models using five algorithms with multiple feature selection strategies, achieving robust predictive performance for DbpA, OspC, and RevA variants (all performance metrics >0.7). Feature importance analysis identified key predictive amino acid residues for DbpA, OspC, and RevA. Notably, B-cell epitope prediction revealed significant enrichment of ML-identified residues within predicted epitope regions for OspC and RevA, suggesting these residues may influence immune recognition and bacterial persistence. This study establishes the first computational framework linking *Borrelia burgdorferi* protein sequence variants to clinical dissemination phenotypes, providing molecular insights into Lyme disease pathogenesis that may inform development of improved diagnostics and therapeutic targets.

## 1. INTRODUCTION

Lyme disease (LD) is an emerging infectious tick-borne disease in the northern hemisphere, with over 470,000 cases diagnosed and treated annually in the United States (1). The causative agents are spirochetes belonging to the *Borrelia burgdorferi sensu lato* complex, collectively referred as Lyme borreliosis (LB). In North America, *B. burgdorferi sensu stricto* (hereafter Bb) predominates. Clinical manifestations range from localized erythema migrans to severe disseminated complications including Lyme arthritis, carditis, and neuroborreliosis, with treatment outcomes and disease severity varying widely among patients (2, 3). Understanding the bacterial determinants that influence disease progression from localized to disseminated infection is crucial for improving diagnostic strategies, treatment protocols, and patient outcomes.

LB exhibits a remarkable level of genetic diversity (4, 5). Its genome consists of an approximately 900 kb linear chromosome and about 21 plasmids (9 circular and 12 linear) ranging from 5 to 84 kb in size (5–7). The genomic background, particularly plasmid content, affects its ecological, epidemiological, and pathogenic properties (8–14). Clinical and animal studies have demonstrated that variations in plasmid content and sequence diversity of virulence factors influence spirochete dissemination and disease severity in mice and humans (15–17). However, the segmented genome and high plasmid variability create technical challenges for sequencing, assembly, and analysis LB genome, ultimately limiting the development of consistent diagnostics and treatments. Since the publication of the first complete Bb genome in 1997 (7), numerous efforts have been made to sequence and map LB genomes (4, 8, 18–20). In 2023, Lemieux et al. published the first large-scale long-read whole genome sequence (WGS) of human Bb isolates (21), providing a valuable resource for examining associations between Bb genotypes and clinical outcomes.

Bb is a Gram-negative bacterium with an atypical cell envelope lacking lipopolysaccharide, but enriched with outer membrane glycolipids and lipoproteins that are highly immunoreactive (22, 23). Many of these lipoproteins function act as virulence factors critical for host-pathogen interactions, including vascular transmigration, immune invasion, and tissue adhesion. By analyzing the WGS of 299 human isolate, Lemieux et al., (2023) demonstrated that surface exposed lipoproteins encoded on a unique set of plasmids and several loci are linked to dissemination. Several studies have also reported the bindings of surface-exposed lipoproteins to the host extracellular matrix (ECM) such as fibronectin (24–26), glycosaminoglycans (GAGs) (27–29), and plasminogen (30–33). Despite the established importance of these surface lipoproteins in pathogenesis, the specific sequence variations that determine their functional differences and contributions to dissemination capacity remain poorly characterized.

Machine learning (ML) for classification tasks has been used for decades in biomedical studies. The application of ML approaches to pathogen genomics has shown remarkable success in identifying novel virulence determinants, predicting antimicrobial resistance, and classifying disease severity across various bacterial pathogens (34–36). These computational methods are adept at detecting subtle patterns in high-dimensional biological data that may elude traditional statistical analyses. Algorithms such as Lasso and elastic-net regularized Generalized Linear Model (GLM), Principal Component Analysis Neural Network (PCANN), Partial Least Square (PLS), Support Vector Machine (SVM), and Random Forest (RF) are well-suited for high-dimensional, small-sample-size datasets, feature selection, and overfitting control (37–42). These tools have also proven their efficiency in supporting biological interpretation and biomarker discovery (43–47).

This study aimed to identify specific amino acid (aa) residues within well-characterized LB surface-exposed virulence factors that could serve as biomarkers for predicting human dissemination phenotypes. We integrated genomic data from a large collection of clinical isolates with advanced computational methods. The surface-exposed proteins examined (BBK32, DbpA, OspA, OspC, P66, and RevA) were selected based on their established roles in host cell adhesion, immune evasion, and tissue invasion processes critical to bacterial dissemination. Using multiple ML algorithms, feature selections, and resampling methods, we successfully developed highly accurate predictive models for DbpA, OspC, and RevA sequences. This computational framework demonstrates the potential of sequence-based virulence prediction and provides molecular insights that could advance LD research.

## 2. MATERIAL AND METHODS

### 2.1 Data collection

Raw whole genome sequence reads and associated metadata of 299 clinically isolated Bb strains were downloaded from NCBI BioProject PRJNA923804. Sequences were trimmed and assembled using Trimmomatic v0.39 (48) and SPAdes 3.15.5 (49). Assembled sequences were annotated based on B31 translated coding sequences using Prokka (50).

Sequences of BB_0406, BBK32, DbpA, OspA, OspC, P66, and RevA from all strains were collected into a single fasta file per protein. Using R packages DECIPHER v2.28.0 (51) and Biostrings v2.68.1 (52), we identified a unique set of sequences for each protein. Sequences were aligned and removed signal sequences prior to feature calculations. To assign names to each sequence, the distance between each sequence and the B31 reference sequence was calculated and ranked from smallest to largest distance. Sequences were then named based on their representative rank. For OspC, sequences were named as the OspC type information from the metadata file. When there is more than one unique sequence mapped to one OspC type, an addition number is added after OspC type to indicate variants in the OspC type.

For each unique sequence, we collected clinical data regarding the strains that possess that sequence. If any of the strains was collected from disseminated Lyme disease patient, the sequence will be labeled as Dis for its dissemination potential. Otherwise, the sequence is labeled as Non-dis. If no dissemination status available for all strains associated with the sequence, its dissemination possibility is NA and the sequence is removed from further analysis. Information of all unique sequences of the seven proteins are listed in Supplementary Table S1.

### 2.2 Cramér’s V association analysis

To assess the strength of association between two categorical variables (protein variants and dissemination phenotype), Cramér’s V analysis was used (53). Cramér’s V is the expansion of the chi-square test for contingency tables bigger than 2 × 2. Cramér’s V values range from 0 (no association) to 1 (perfect association). A value bigger than 0.25 is named as a very strong relationship (54).

### 2.3 Machine learning modeling of dissemination potential

#### 2.3.1 Data preprocessing

For each protein, unique protein sequences were aligned, and signal sequences were removed. Each sequence was transformed to a binary feature matrix using one-hot encoding, a standard orthogonal encoding technique for aa sequences (55). Each individual sequence matrix has dimensions L × D, where L represents the sequence length and D = 21 (20 standard aa plus 1 gap character “-“ from the alignment). The individual matrices were concatenated to form a single feature matrix of shape (N, L × D), where N is the total number of sequences. To reduce dimensionality and eliminate uninformative features, columns with constant values across all sequences were removed. The resulting matrix was used as the input feature matrix for ML modeling.

#### 2.3.2 Feature selection

Feature selection is a widely used strategy in machine learning to select a subset of features from the original set, aiming to optimize learning performance while reducing computational expense (56, 57). In this study, we employed both supervised and unsupervised feature selection approaches to generate seven different feature sets for modeling each protein.

**Supervised feature selection:** Features were ranked using model-independent Variable Importance (VIP) scores calculated as the Area Under the Receiver Operating Characteristic Curve (AUC-ROC) for each individual feature. A VIP score of 0.5 indicates no discriminative power. Four subsets of features were created: (1) fs055 - features with VIP ≥ 0.55, and (2–4) ImpQ1, ImpQ2, and ImpQ3 - features with VIP scores above the first, second, and third quartiles, respectively.

**Unsupervised feature selection**: Features were selected based on variance filtering. Three feature sets (VarQ1, VarQ2, and VarQ3) were created using the first, second, and third quartiles of features variance as selection thresholds. The numbers of features in the original data and each subset are listed in Supplementary Table. S2.

#### 2.3.3 Model training and class imbalance handling

The dataset presented two main challenges: small sample size and class imbalance with the disseminating class (Dis) containing about twice the number of sequences in the non-disseminating class (Non-dis). To address class imbalance, we applied Synthetic Minority Oversampling Technique (SMOTE) to oversample the minority Non-dis class (58).

For robust model evaluation, we employed stratified sampling with 25 repetitions of 4-fold cross-validation, generating 100 distinct train-test splits. Five ML algorithms were used: Partial Least Square Discriminant Analysis (PLS), Random Forest (RF), Support Vector Machine (SVM), Generalized Linear Model (GLM), and Principal Component Analysis Neuron Network (PCANN). Two resampling methods Leave-One-Out Cross Validation (LOOCV) and bootstrap were utilized to tune hyperparameters.

#### 2.3.4 Model performance evaluation

The performance of models was evaluated using multiple metrics: AUC-ROC, accuracy, sensitivity, and specificity. The accuracy, sensitivity, or recall, which is also known as true positive rate (TPR), and specificity which is equal to 1-false positive rate (FPR), were calculated from a confusion matrix using a decision threshold of 0.5. The ROC curve is generated by plotting the TPR against the FPR at various threshold settings. All the performance metrics were computed for both train and test sets to assess potential overfitting and model generalization.

#### 2.3.5 Feature importance analysis

Feature importance was estimated using a model-based approach implemented through the variable importance functions (varimp) in the R ‘caret’ package (59). For each predictive algorithm, features were ranked by their VIP scores, and the top 20 features were selected. The final set of important features for each protein was determined by taking the union of: (1) the intersection of top features across all supervised feature selection models (fs055, ImpQ1, ImpQ2, ImpQ3), and (2) the intersection of top features across all unsupervised feature selection models (VarQ1, VarQ2, VarQ3).

All data preprocessing, model training, and evaluation were conducted using the ‘caret’ package version 6.0 on R version 4.3.2 (59, 60).

### 2.4 B-cell epitope prediction

To predict conformational B-cell epitopes in Bb proteins, we employed the DiscoTope 3.0, a robust structure-based epitope prediction tool that integrates inverse folding latent representations and a positive-unlabeled learning strategy (61). DiscoTope 3.0 has been benchmarked to maintain high predictive performance across solved, relaxed, and predicted structures and outperformed other prediction tools in predicted structures.

3D protein structures of DbpA, OspC, and RevA were generated using AlphaFold 3.0 server (62) based on the protein sequences in B31 strain. The protein structures were submitted to the DiscoTope 3.0 server (https://services.healthtech.dtu.dk/services/DiscoTope-3.0/), which was run using default parameters.

DiscoTope 3.0 computes an epitope propensity score for each surface-exposed residue, based on updated statistics from a larger set of antigen-antibody complex structures. Residues with a calibrated score above the recommended threshold (default: 0.9, recall up to ∼50 % for moderate confidence) were classified as part of a predicted conformational B cell epitope. All predicted residues were visualized and mapped onto the protein structure using Chimera X (63) for spatial localization and further interpretation.

### 2.5 Host-pathogen protein-protein interaction (PPI) site prediction

We computed protein-protein complex structures of DbpA-decorin, OspC-OspC-plasminogen, and RevA-fibronectin using AlphaFold 3 server. For DbpA, OspC, and RevA, aa sequences in B31 strains were used and signal peptides were removed. Sequences of human decorin, plasminogen, and fibronectin were accessed from NCBI NP_001911.1, AAA60113.1, and NP_997647.2, respectively. The sequences of leucine-rich repeat regions in decorin (residues 52-359), five kringle domains in plasminogen (residues 99-570), and 70 kDa regions in fibronectin (residues 52-608), which are known interaction sites with DbpA (64), OspC (65), and RevA (24), respectively, were used to generate the protein-protein structures. Best predicted structures were submitted to Chimera X for visualization and spatial localization.

## 3. RESULTS

### 3.1 Associations of protein variants and dissemination phenotype

We extracted gene sequences of 7 known Bb virulence factors from the published WGS of 299 clinical isolates (NCBI BioProject PRJNA923804). Following translation into aa sequences, we identified between 4 and 36 unique variants per protein. The distribution of these variants with respect to clinical dissemination status is presented in Fig. 1. Most variants were observed in both disseminated and localized (non-disseminated) strains. However, a subset of variants appeared to be exclusively associated with one clinical outcome. Specifically, 2 of 34 variants of BBK32 variants (BBK32_2 and BBK32_28), 1 of 30 variants of DbpA (DbpA_15), 1 of 33 variants of OspC (S_2), 2 of 25 variants of P66 (P66_6 and P66_25), and 4 of 36 variants of RevA (RevA_3, 22, 23, and 24) were found only in strains from disseminated infections. Conversely, 1 of 4 variants of BB_0406, 14 of 34 variants of BBK32, 7 of 30 variants of DbpA, 3 of 15 variants of OspA, 9 of 33 variants of OspC, 13 of 25 variants of P66, and 9 of 36 variants of RevA were found only in localized infections. These findings suggest some protein variants may be associated with either localized or disseminated disease, supporting their potential relevance in virulence and host-pathogen interaction dynamics.

**Figure 1.**
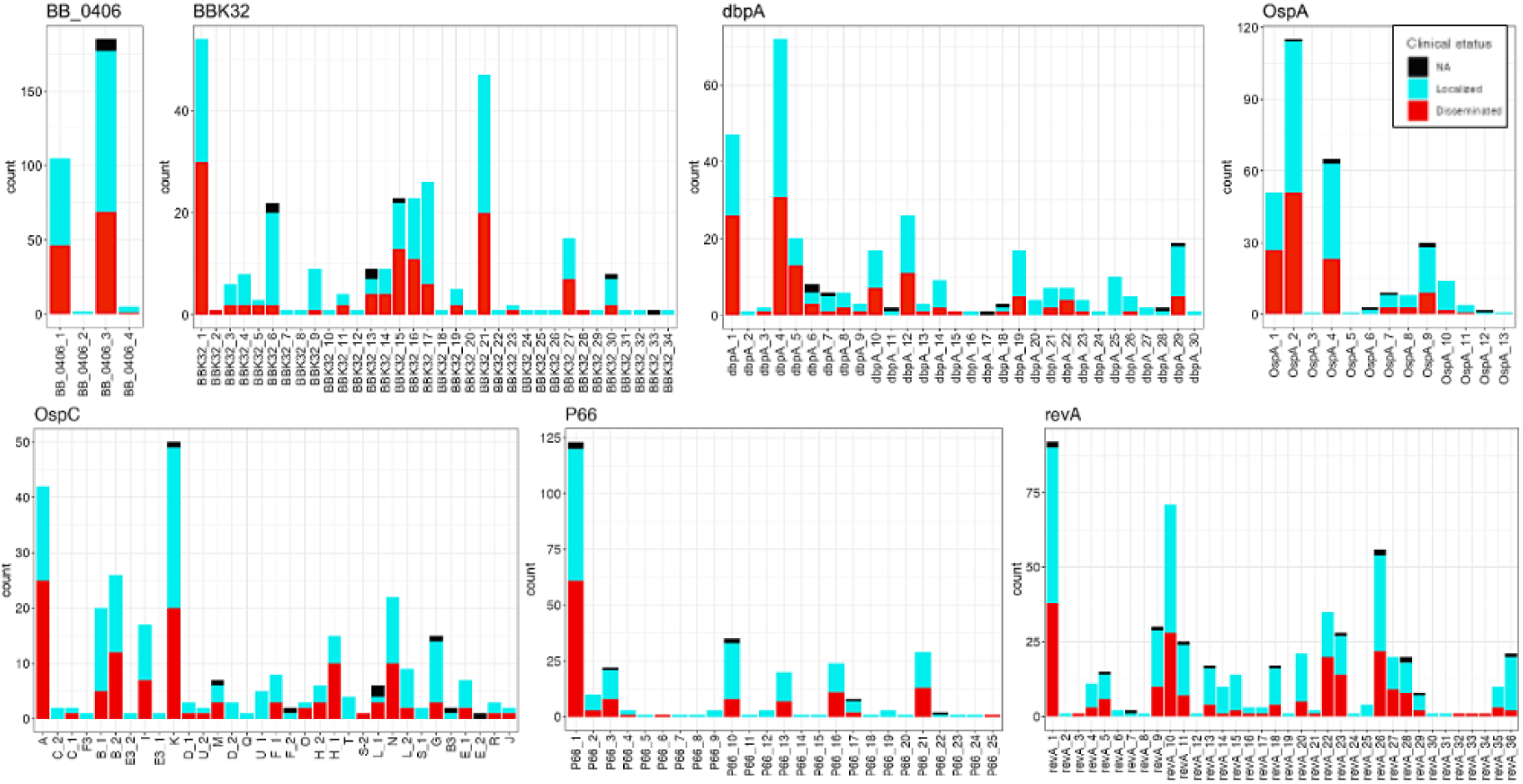
Profile of unique protein sequences of BB_0406, BBK32, DbpA, OspA, OspC, P66, and RevA. Each bar represents the number of isolates carrying the corresponded sequences. All variants are sorted based on their sequence distance from the B31 reference sequence.

**Figure 2.**
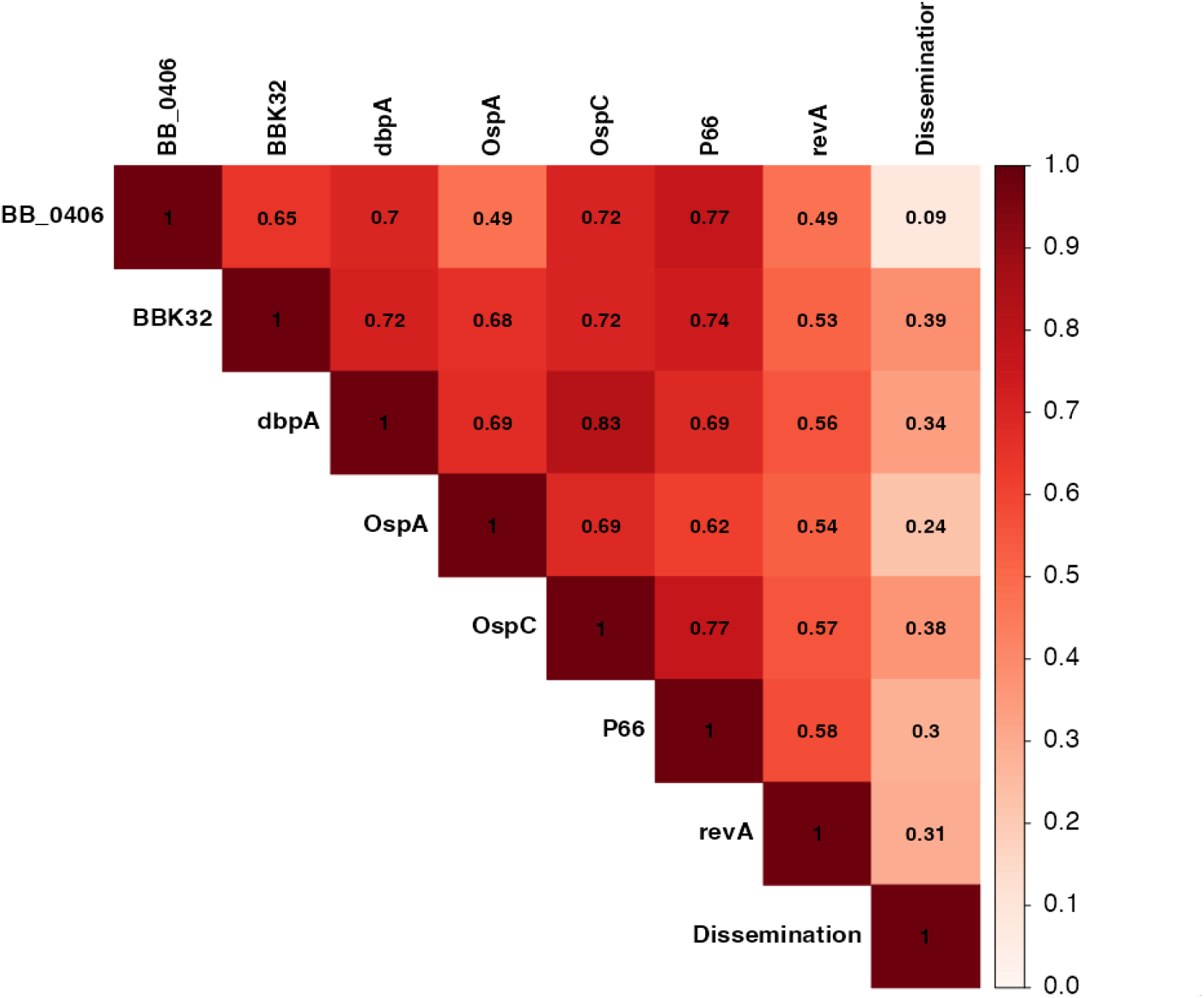
Cramer’s V correlation heatmap. Color intensity reflects the strength of association between variables. Cramér’s V close to 0 indicates no association, >0.10 denotes moderate, and >0.25 denotes strong association (Akoglu, 2018).

To assess the strength of association between protein variants and clinical dissemination status, we performed Cramér’s V analysis. This revealed strong interdependence among all seven proteins, but only BBK32, DbpA, OspC, P66, and RevA showed strong associations with dissemination (Cramér’s V >0.25, (54)). Based on these findings, we selected these five proteins for further ML modeling to classify variants as associated with dissemination or localized infections.

### 3.2 Prediction ability of trained models

We developed 70 different prediction models for each of five proteins (BBK32, DbpA, OspC, P66, and RevA) by combining 7 feature sets, 2 resampling methods, and 5 ML algorithms. Models built using variant sequences from DbpA, OspC, and RevA demonstrated strong predictive performance across both train and test sets with AUC-ROC values, accuracy, sensitivity, and specificity all exceeding 0.7 (95% CI widths < 0.05) (Fig. 3). BBK32 and P66 models showed weaker performance with AUC-ROC values around 0.6 and exhibited high variability in test sets. Comprehensive performance metrics of all models are provided in Table S3. While certain models significantly outperformed others, no single model type consistently achieved optimal results across all proteins (Supplementary Fig. S1-S5).

**Figure 3.**
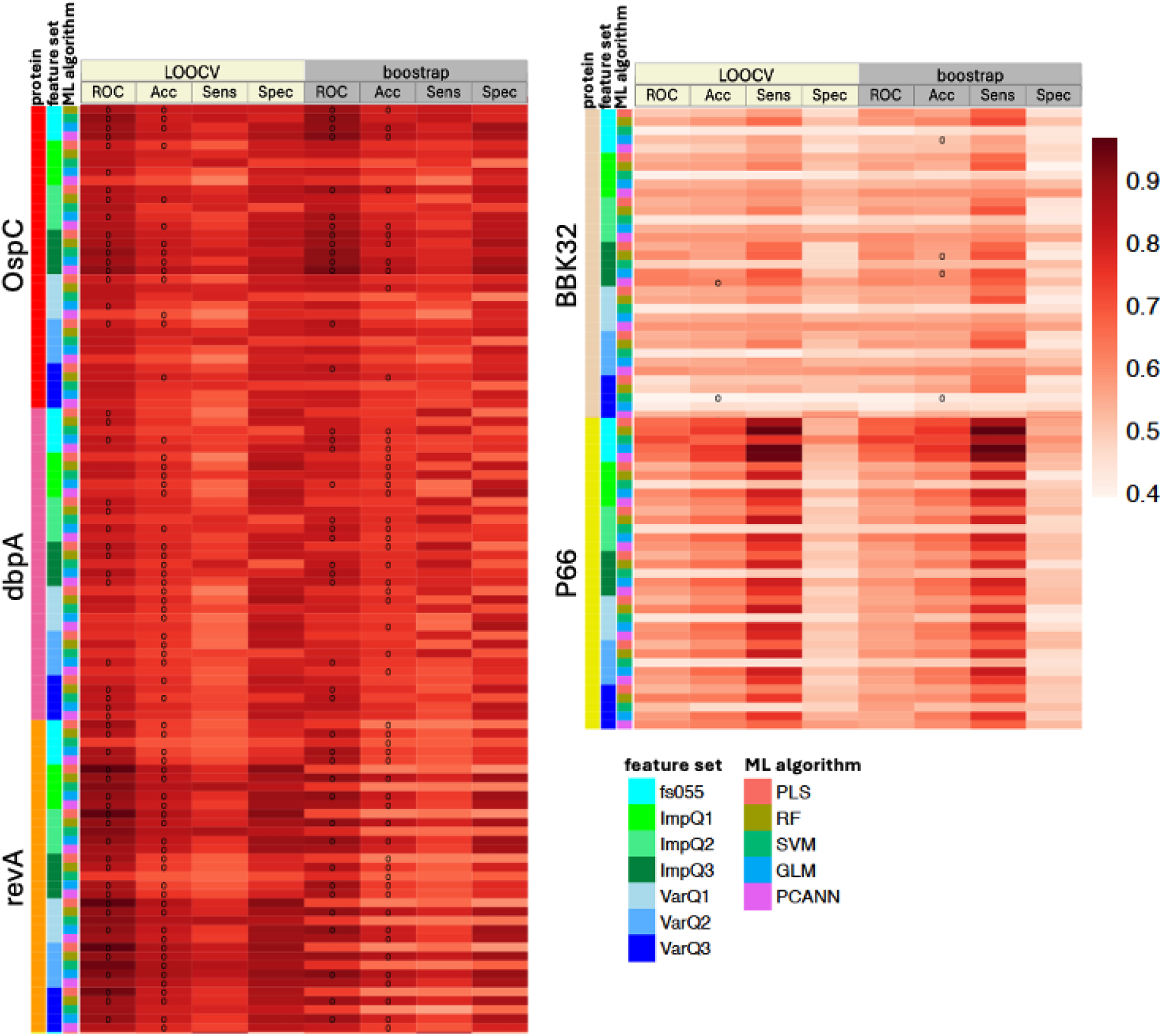
Model performance comparison across different algorithms and metrics. The heatmap displays average performance values for four key metrics evaluated on test datasets: Area Under the ROC Curve (ROC), accuracy (Acc), sensitivity (Sens), and specificity (Spec). Each cell represents the mean performance value for a specific model-metric combination. Black circles within cells indicate statistically significant results where the 95% CI width is less than 0.05. See color scale for details.

To access the impact of different resampling methods on model performance, stability, and generalization, we compare the performance values, coefficient of variation, and train-test differences in LOOCV and bootstrap models. We showed that LOOCV consistently generated superior performance metrics across all feature sets, algorithms, and proteins, with significantly higher AUC-ROC values, accuracy, and specificity compared to bootstrap validation (Supplementary Fig. S6). The LOOCV models demonstrated substantially greater model stability, as evidenced by narrower confidence intervals and reduced variability in performance metrics across the 100 train-test splits (Supplementary Fig. S7). The train-test performance differences were also significantly smaller in LOOCV models indicating more consistent generalization and potentially reduced sensitivity to data variation or regularization effects (Supplementary Fig. S8). Additionally, the Bland-Altman and correlation analyses showed strong overall agreement between LOOCV and bootstrap methods across all metrics (ROC, Accuracy, Sensitivity, and Specificity), with Pearson’s r values ranging from 0.81 to 0.93 (Supplementary Fig. S9-10). However, some variability and wider limits of agreement were observed for certain proteins, particularly RevA. This suggests that while the two methods largely produce consistent results, a few cases may be more sensitive to resampling techniques.

### 3.3 Model-based important features

We identified top 20 features in each model and then determined the set of important features or key aa residues for DbpA, OspC, and RevA as described in the Materials and Methods section. There were 57, 29, and 42 features were selected for DbpA, OspC, and RevA, respectively. Fig. 4 shows the frequency of the key aa residues in Dis or Non-dis sequences. Several of these residues display distinct prevalence patterns between the two groups, suggesting their role in differentiating dissemination potential.

**Figure 4.**
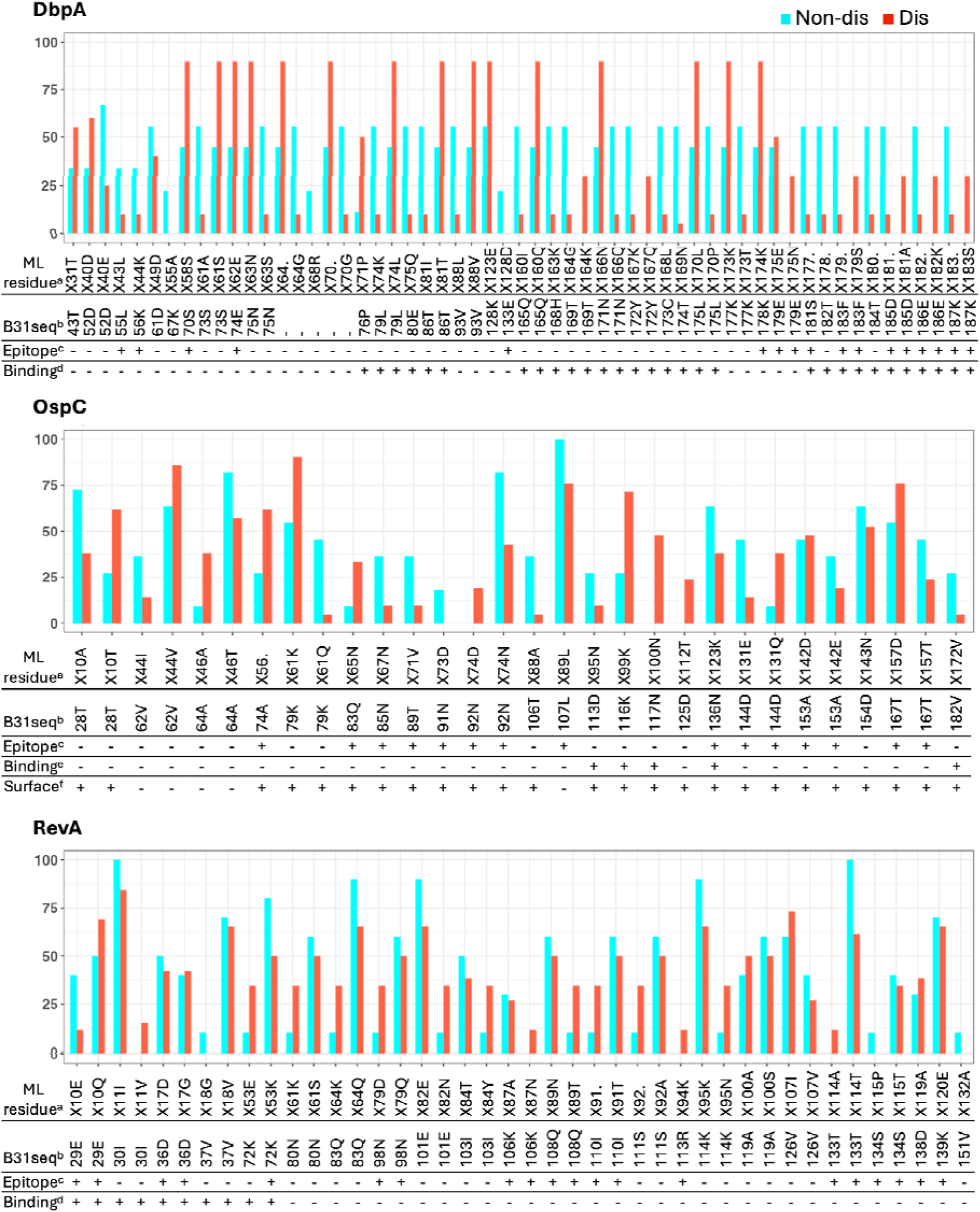
Prevalence of important features in dissemination (Dis) and non-dissemination (Non-dis)-linked variants of DbpA, OspC, and RevA. **^a^** Residue labels as defined in machine learning models. ^b^ Corresponding amino acid residues in the B31 reference sequence. ^c^ Residues predicted to be part of B-cell epitopes by DiscoTope 3.0. ^d^ Residues previously reported to be important for binding to host proteins (24, 93–95). ^e^ Residues located at the interface of the plasminogen-OspC-OspC complex, as predicted by AlphaFold3 and ChimeraX. ^f^ Residues with solvent-accessible surface area > 20 Å² in the OspC dimer as calculated using Chimera X.

### 3.4 B-cell epitope prediction

To assess the potential immunological relevance of the predictive residues, we predicted B-cell epitopes in the DbpA, OspC, and RevA variants of B31 strain using AlphaFold3 and DiscoTope 3.0. The DiscoTope scores for individual residues in each proteins are listed in Table S4-S6. DiscoTope identified 35, 48, and 49 residues within predicted B-cell epitope regions in DbpA, OspC, and RevA, respectively, using a moderate confidence threshold (0.9, recall up to 50%). We found some overlaps between the predicted epitope residues and ML-based predictive residues (Fig. 4).

To evaluate whether the observed overlaps exceeded random expectation, we performed one-side Fisher’s Exact test for each protein. For OspC, the overlap was 11 residues (OR = 3.57, *p* = 0.006), suggesting significant enrichment of dissemination predictive residues within predicted epitopes. RevA also exhibited significant enrichment (12 overlapping residues, OR = 2.37, *p* = 0.048). DbpA also had 12 overlapping residues with a potential association (OR = 2.12), although the result was not statistically significant (*p* = 0.058). These results support a biologically meaningful convergence between structural epitope prediction and ML-based residue importance.

### 3.5 Host-pathogen PPI site prediction

DbpA, OspC, and RevA have been well-known to bind to decorin, plasminogen, and fibronectin, respectively. We hypothesized that some of the predictive residues might be located at the interfaces of these PPIs. To explore this, we generated structural models of decorin-DbpA, plasminogen (5-kringle domain)-OspC-OspC, and fibronectin (70 kDa region)-RevA complexes using AlphaFold 3. The predicted TM scores (pTM) for the decorin-DbpA and plasminogen-OspC-OspC complexes exceeded 0.5, suggesting that the overall folds may reasonably accurate. However, the pTM of the fibronectin-RevA complex was only 0.3, and the interfacial predicted TM scores (ipTM) - which assess the relative positioning accuracy between subunits - were low for both decorin-DbpA (0.17) and fibronectin-RevA (0.11) complexes, indicating unreliable interface predictions. The plasminogen-OspC-OspC complex had a moderate ipTM of 0.53, suggesting borderline confidence. Given these limitations, the predicted complex structures were not fully reliable for detailed interface analysis. Nevertheless, mapping the ML-identified predictive residues onto the plasminogen-OspC-OspC model revealed that some may indeed lie within potential interaction sites (Supplementary Fig. S11B). In addition, we generated OspC dimer independently, which showed a high structural similarity to the published crystal structure (PDB: 1GGQ), with an RMSD of 0.257 Å (Supplementary Fig. S11A). The predicted OspC dimer achieved both pTM and ipTM of 0.83. Most of the predictive peptides locate on the accessible surface area of the dimer (Fig. 4 and Supplemental Fig. S11A).

## 4. DISCUSSION

This study represents the first comprehensive ML-based analysis of Bb protein variants and their association with clinical dissemination phenotypes. Through systematic examination of seven known virulence factors across 299 clinical isolates, we identified specific protein variants and aa residues that demonstrate strong predictive capacity for distinguishing between disseminated and localized infections. These findings provide novel insights into the molecular determinants of LD pathogenesis and potential biomarkers for clinical prognosis.

## Key findings and their implications

By leveraging WGS data from 299 clinical Bb isolates and employing a combination of correlation analysis and ML models, we demonstrated that variations in several adhesins, particularly DbpA, OspC, and RevA, are significantly associated with the clinical phenotype of dissemination. Our findings underscore the importance of surface-exposed adhesins in the pathogenesis of LD and provide a molecular framework for their continued investigation as potential targets for diagnostics and therapeutics.

Surface-exposed adhesins are integral to Bb’s ability to colonize host tissues, evade immune defenses, and disseminate throughout the host (66, 67). Among the adhesins analyzed, five proteins - BBK32, DbpA, OspC, P66, and RevA-demonstrated significantly strong associations with dissemination phenotypes. The lack of strong association between BB_0406 and OspA in Cramér’s V analysis may be partially attributed to the small number of variants observed for these proteins.

BBK32 is a fibronectin- and GAG-binding adhesin essential for vascular interactions and joint colonization (25, 26, 68, 69). RevA, another fibronectin-binding protein with elevated expression during mammalian infection, has been shown to influence dissemination, arthritis severity, and host response (24, 70, 71). DbpA, a decorin binding protein that also possesses GAG-binding activity, plays a critical role in tissue colonization and early infection establishment (72–74). P66, functioning as both an integrin adhesin and porin, has been implicated in endothelial transmigration and tissue invasion (75, 76). OspC is another adhesin crucial for early-stage infection with a dual role in host cell adhesion and evasion of the complement system, exhibiting substantial sequence diversity across Bb strains (77–80). Certain OspC types are preferentially associated with invasive phenotypes (10, 81–83). Comparative analysis of BBK32 orthologues from different LB species reveals that both *B. garinii* and *B. afzelii* variants bind C1 and C1r complement components with high affinity similar to Bb BBK32 though only the *B. afzelii* BBK32 orthologue (BGD19) demonstrates potent inhibition of the classical complement pathway like Bb BBK32 (84). Similarly, DbpA variants across different LB species have been shown to affect spirochetes binding to decorin, dermatan sulfate, and mammalian cells, as well as influence tissue tropism in mice (85, 86). Our study showed that within Bb, there are certain variants of these proteins exclusively associated with only non-invasive or invasive infections in humans. The results further support potential roles of these proteins in tissue invasion and provide essential reference points for studying interspecies variations in pathogenesis mechanisms and clinical outcomes.

We successfully built robust ML models capable of distinguishing protein sequences of DbpA, OspC, and RevA that are potentially associated with dissemination, with all performance metrics exceeding 0.7. This represents a novel computational approach to predicting Bb virulence potential based solely on protein sequence data. Our findings reveal that specific aa residues within these surface-exposed proteins serve as critical determinants of dissemination capacity. The identification of 57, 29, and 42 key predictive residues for DbpA, OspC, and RevA, respectively, provides molecular-level insights into the mechanisms underlying Bb pathogenesis.

## Biological relevance of predictive peptides

The convergence between our ML-identified predictive residues and computationally predicted B-cell epitopes provides compelling evidence for the biological relevance of our findings. The significant enrichment of predictive residues within predicted epitope regions for OspC (OR = 3.57, *p* = 0.006) and RevA (OR = 2.37, *p* = 0.048) suggests that immune recognition of these specific residues may play a crucial role in determining disease progression. Variants with epitope residues that are less immunogenic or exhibit altered antigenicity may have enhanced capacity to disseminate by avoiding rapid immune clearance.

These results are particularly noteworthy when compared to previously characterized epitopes in these proteins. The Immune Epitope Database (IEDB) (www.iedb.org) is a comprehensive resource that catalogs experimentally validated immune epitopes from infectious, allergic, and autoimmune disease and transplant rejection studies in human, non-human primates and other animal species (87). Many of our predicted predictive and epitope residues of OspC and DbpA (no data available for RevA) lie within the linear B-cell epitopes extracted from IEDB (Supplemental Fig. S14). A previous study reported that the human B-cell epitopes regions on OspC, including residues 71-86, 104-118, 156-171, and 184-190, are within the binding sites of OspC and mouse B5 antibody (88). B5 is the only mouse OspC-specific monoclonal IgG2a antibody (MAb) that has been demonstrated to protect mice against Bb challenge via either needle injection and tick infection (89–91). Interestingly, the OspC predictive residues in the residues 71-86 and 104-118 were observed mainly in non-dissemniated isolates (Fig. 4). Collectively, these findings underscore the potential role of immune-targeted residues in influencing pathogen dissemination, reinforcing the relevance of our predictive approach to identifying biologically meaningful targets.

We hypothesized that predictive residues play roles in host-pathogen interactions. Unfortunately, all the host-pathogen protein complex structures predicted by AlphaFold3 had very low ipTM values, which implied failed prediction in the relative orientation of interacting partners. Although this limits the identification of specific interface residues for each complex, we observed some of the OspC predictive residues located within potential interaction sites of plasminogen-OspC-OspC (Supplemental Fig. S11B). In addition, although the dimer interface of OspC buried approximately 22% of the accessible surface area of each unit (92), most of the OspC predictive residues were located outside this buried region (Supplemental Fig. S11A). This result indicates their potential availability for interactions with host proteins and/or recognition by the immune system, suggesting potential role in disease dissemination.

For DbpA, several residues which include Lysines 82, 163, 170 and the C-terminal tail have been characterized to be important for GAG bindings (93, 94). The three Lysines are conserved in all DbpA variants in our study. The peptide sequence 76-90 was shown to be crucial for decorin-binding while peptide sequence 152-176 retains the ligand binding activity by maintaining the appropriate decorin-binding conformation (95). While these experimentally validated peptide regions did not overlap with the binding site in our predicted decorin-DbpA structure - likely due to the small ipTM value - several of our ML-based predictive residues are located within these regions and the C-terminal tail (Fig.4 and Supplementary Fig. S12). A similar pattern was observed with RevA, where the first 60 aa residues on the N-terminus, previously shown to be essential for fibronectin binding (24), did not overlap with the binding site on the predicted fibronectin-RevA structure (Fig. 4 and Supplementary Fig. S13). Nevertheless, out ML models identified several N-terminal residues as important for distinguishing potentially disseminated variants. Collectively, these results support the biological relevance of our predictive residues and their potential roles in mediating host-pathogen interactions crucial for disease dissemination.

## Methodological considerations and model performance

Our comparative analysis revealed that LOOCV resampling method consistently outperformed bootstrap method in model performance, stability, and generalization capability. These findings demonstrate the importance of resampling strategy selection in small-dataset machine learning applications. LOOCV’s superior performance likely stems from the method’s ability to utilize nearly all available data for training while ensuring each sample is tested exactly once (96). This is particularly advantageous when working with limited protein variants where maximizing the use of available data is crucial for robust model development. The bootstrap method’s poorer performance may be attributed to sampling with replacement, which can lead to some samples being over-represented in training sets while others are excluded, potentially creating artificial patterns or missing important variant information (97). This sampling bias becomes more pronounced in small datasets where each unique sequence carries significant informational value. The smaller train-test performance differences in LOOCV models indicated reduced overfitting and more realistic performance estimates. We should note that the slight negative gap (approximately −0.05) in bootstrap models may reflect regularization effects or data variability rather than overfitting. In addition, despite strong overall agreement between methods (Pearson’s r = 0.81-0.93), the wider limits of agreement for certain proteins like RevA highlight protein-specific sensitivities to resampling choice. For small variant datasets, LOOCV should be preferred over bootstrap methods due to superior performance estimation and stability, while bootstrap may be advantageous for larger datasets where computational efficiency is paramount. These findings emphasize that resampling method choice can significantly impact the clinical utility of ML models in infectious disease research.

The weaker performance of BBK32 and P66 models compared to DbpA, OspC, and RevA models may reflect several factors: (1) these proteins may have less direct involvement in dissemination mechanisms, (2) their functional variants may be more context-dependent, requiring consideration of additional factors not captured in our current analysis, or (3) the sample size may be insufficient to detect subtle but biologically relevant associations for these proteins.

## Study limitation and future directions

While our computational approach offers valuable insights, several limitations must be acknowledged. First, the relatively small dataset size and class imbalance required the use of SMOTE oversampling, which may introduce artificial patterns in the data. Second, we simplified the classification of variants into just two classes based on any present/absence of the variants in disseminated isolates. This binary classification approach may oversimplify dissemination as a discrete trait when it likely exists on a continuous spectrum, and sequences labeled as “Non-dis” may reflect incomplete clinical sampling rather than true lack of dissemination capacity. Third, while our structural predictions provide valuable insights, the low ipTM scores for the protein complexes limit the reliability of interface analyses. Future studies incorporating experimental validation of predicted interaction sites through mutagenesis binding assays, and infection models will be necessary to confirm the functional impacts of identified residues. Fourth, our analysis focused on known virulence factors, potentially missing important determinants encoded elsewhere in the Bb genome. Expanding the analysis to include additional surface proteins and exploring whole-genome approaches could reveal novel dissemination-associated factors. In addition, the clinical metadata used to classify dissemination may vary in diagnostic criteria over the 30-year isolate collection period, introducing potential confounding factors.

## Conclusions

This study demonstrates the power of ML approaches in identifying biologically relevant patterns in pathogen genomic data. The strong predictive performance of DbpA, OspC, and RevA variant-based models, combined with the immunological relevance of predictive residues, provides a foundation for both mechanistic understanding and clinical application. The methodological framework developed here may also be applicable to other bacterial pathogens where strain-level variation influences disease outcomes.

## Acknowledgements

The work was supported by the National Institute of Allergy and Infectious Diseases at the National Institutes of Health [grant number 1R01AI158304]. This work used advanced cyberinfrastructure resources provided by the University of North Dakota Computational Research Center.

Molecular graphics and analyses performed with UCSF ChimeraX, developed by the Resource for Biocomputing, Visualization, and Informatics at the University of California, San Francisco, with support from National Institutes of Health R01-GM129325 and the Office of Cyber Infrastructure and Computational Biology, National Institute of Allergy and Infectious Diseases.

## Supplementary Materials

Supplementary Tables (S1-S6) and Figures (S1-S14) are available.

## Data Availability

Code for the study pipeline is available on GitHub: https://github.com/hoa1279/HN_Lyme_ML_2025

