## Supplemental Figure for "Machine Learning-Driven Identification of Virulence Determinants in *Borrelia burgdorferi* Associated with Human Dissemination"

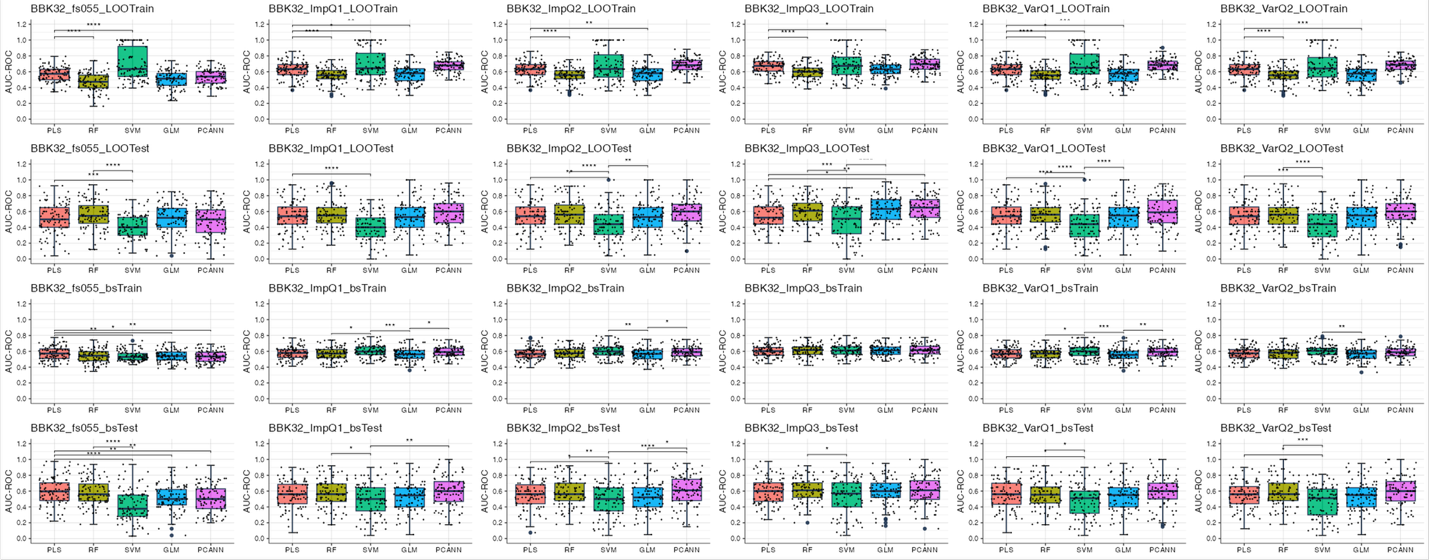

Figure S1: AUC-ROC comparison of machine learning algorithms for BBK32 models in different resampling methods (LOOCV (LOO) and bootstrap(bs)) and different feature sets. Statistical significance of pairwise t-test comparisons between algorithms is indicated above each panel. ****p < 0.0001, ***p < 0.001, **p < 0.01, *p < 0.05.

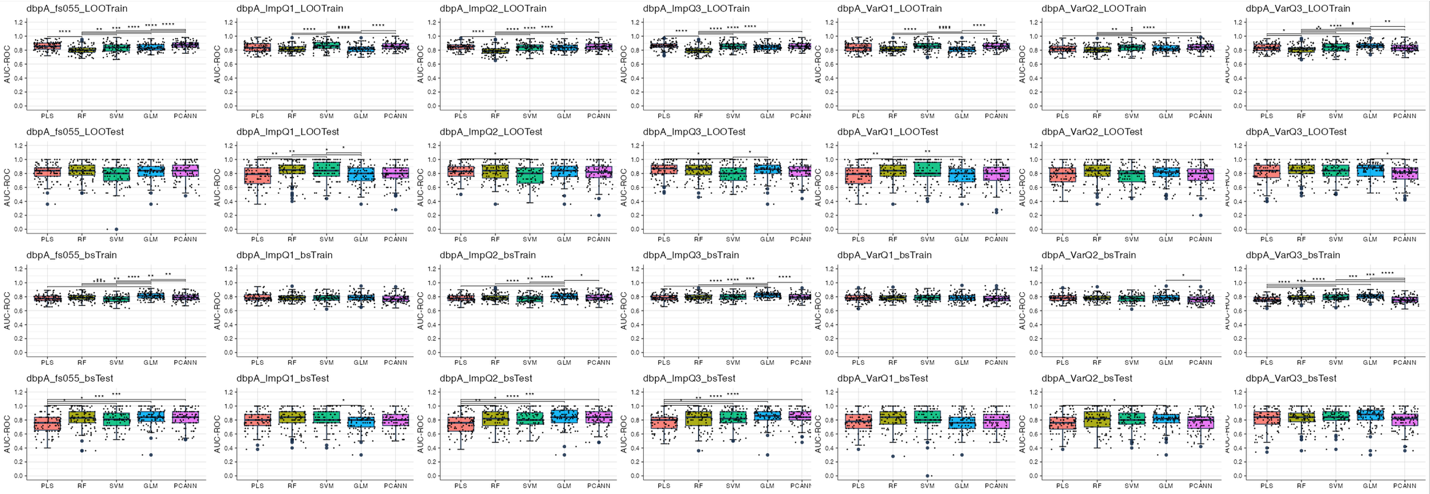

Figure S2: AUC-ROC comparison of machine learning algorithms for DbpA models in different resampling methods (LOOCV (LOO) and bootstrap(bs)) and different feature sets. Statistical significance of pairwise comparisons between algorithms is indicated above each panel. ****p < 0.0001, ***p < 0.001, **p < 0.01, *p < 0.05.

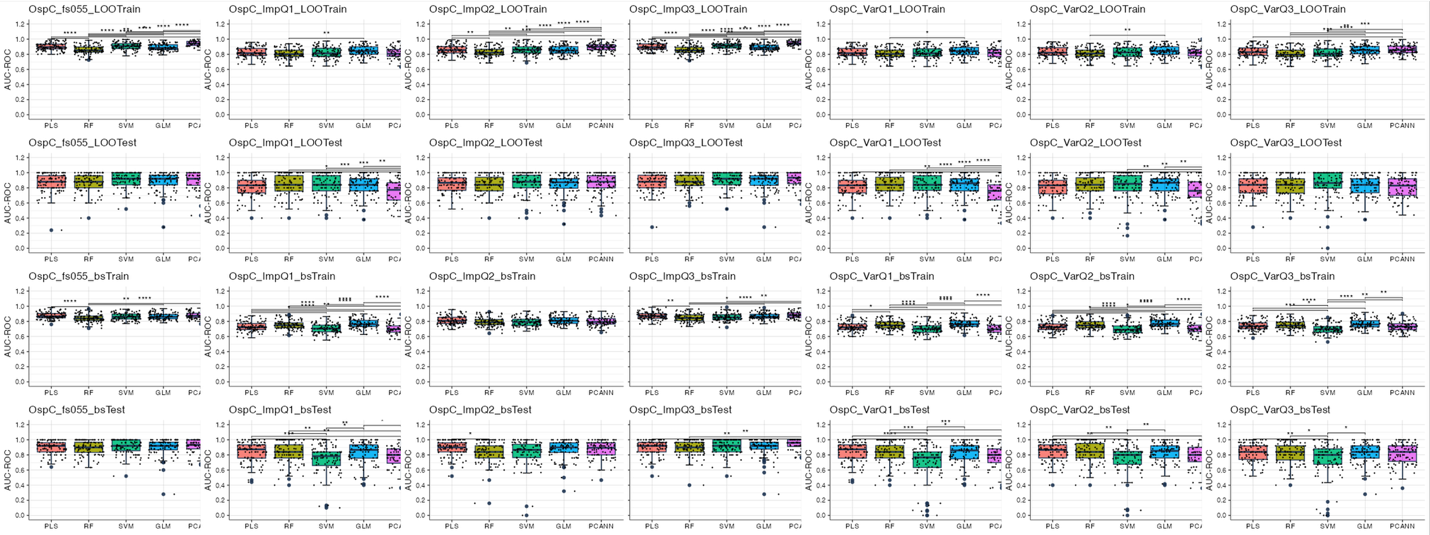

Figure S3: AUC-ROC comparison of machine learning algorithms for OspC models in different resampling methods (LOOCV (LOO) and bootstrap(bs)) and different feature sets. Statistical significance of pairwise t-test comparisons between algorithms is indicated above each panel. ****p < 0.0001, ***p < 0.001, **p < 0.01, *p < 0.05.

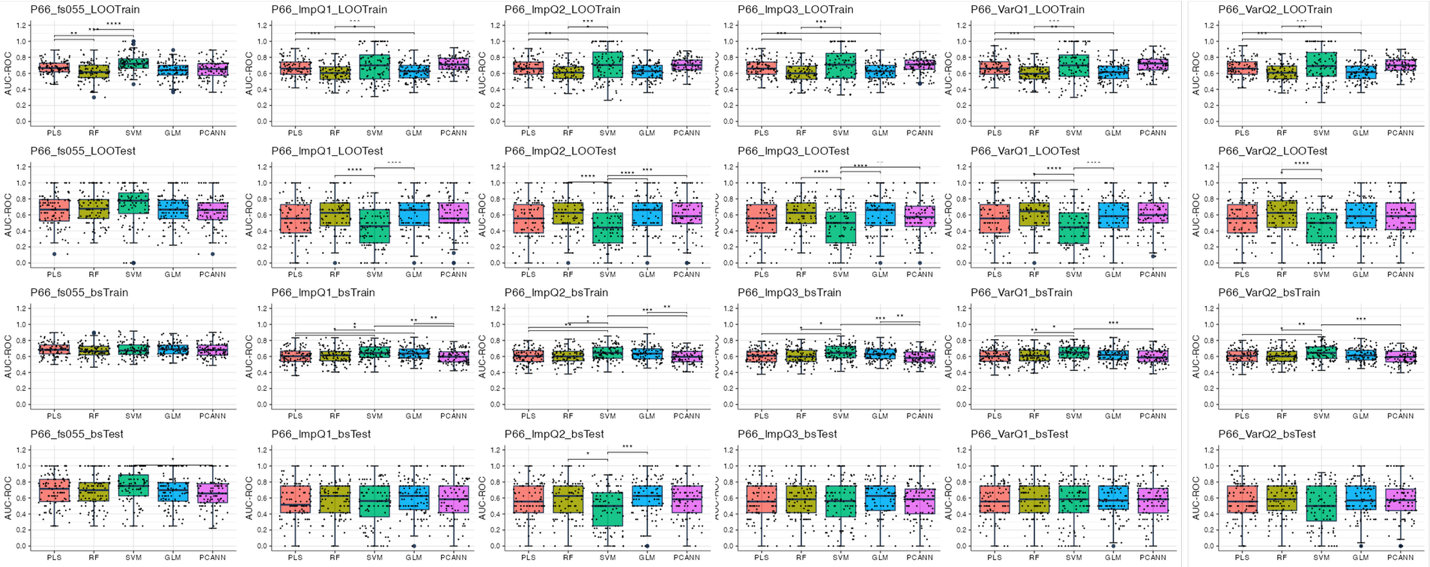

Figure S4: AUC-ROC comparison of machine learning algorithms for P66 models in different resampling methods (LOOCV(LOO) and bootstrap(bs)) and different feature sets. Statistical significance of t-test pairwise comparisons between algorithms is indicated above each panel. ****p < 0.0001, ***p < 0.001, **p < 0.01, *p < 0.05.

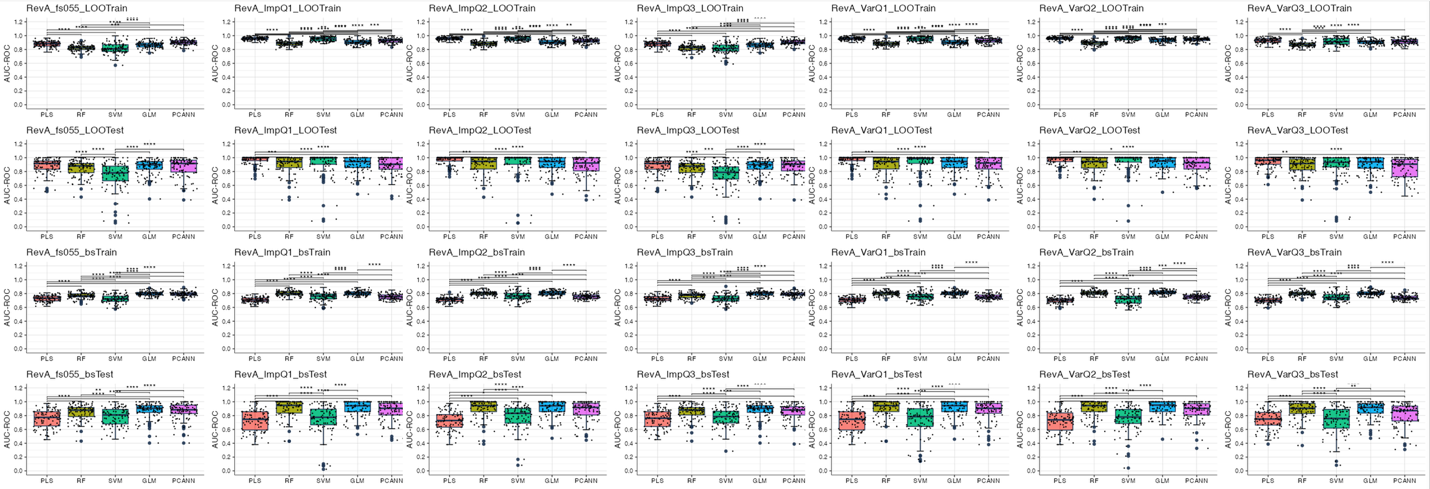

Figure S5: AUC-ROC comparison of machine learning algorithms for RevA models in different resampling methods (LOOCV(LOO) and bootstrap(bs)) and different feature sets. Statistical significance of pairwise t-test comparisons between algorithms is indicated above each panel. ****p < 0.0001, ***p < 0.001, **p < 0.01, *p < 0.05.

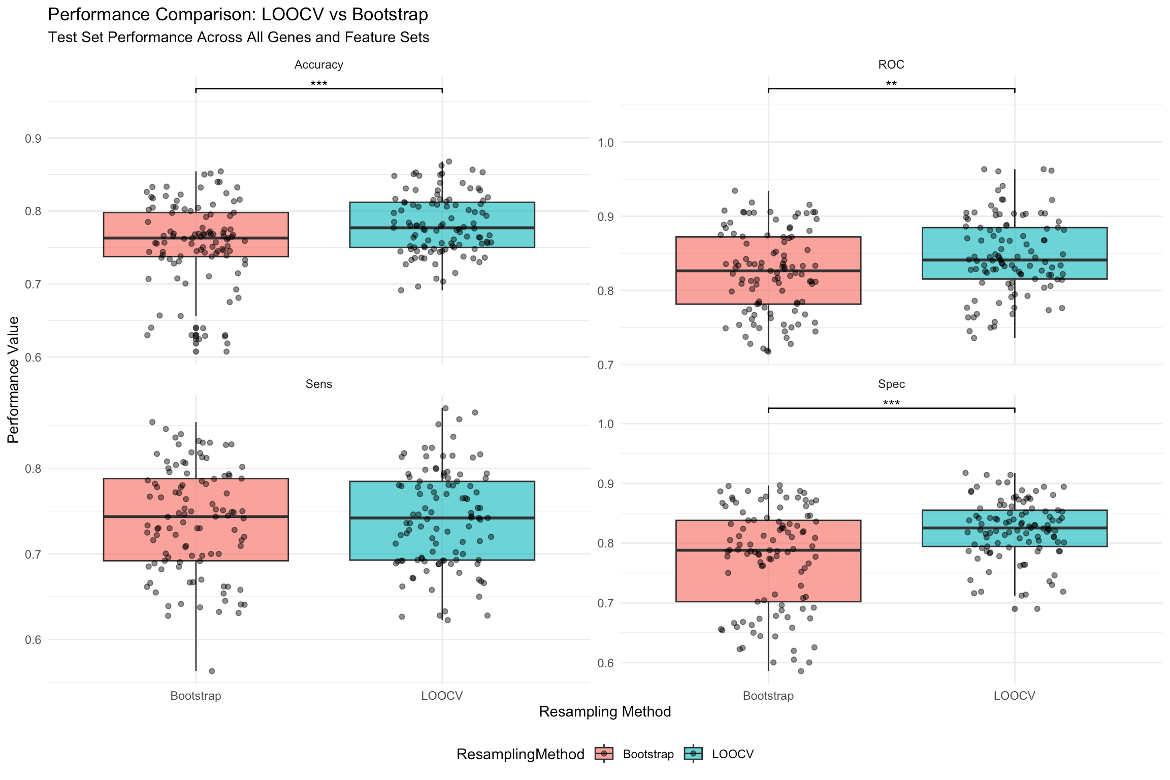

Figure S6: Performance comparison between LOOCV and bootstrap models across all proteins and feature sets. ***p < 0.001, **p < 0.01.

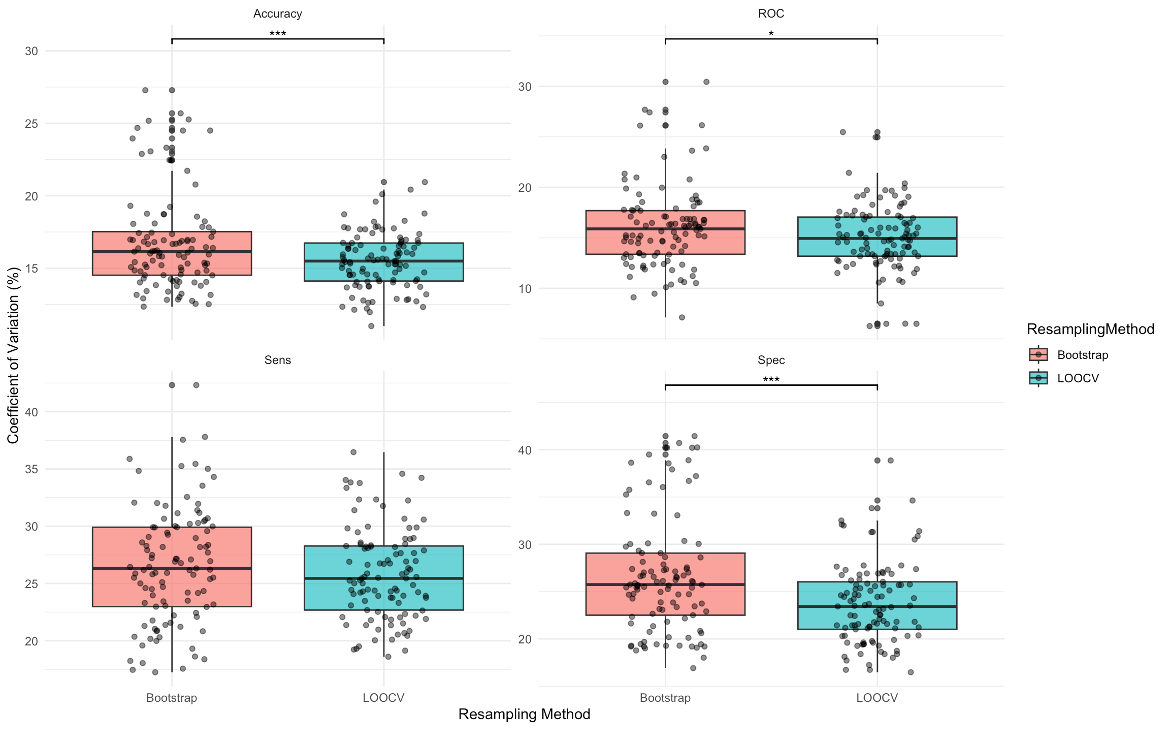

Figure S7: Model stability comparison between LOOCV and bootstrap models across all proteins and feature sets. ***p < 0.001, *p < 0.05.

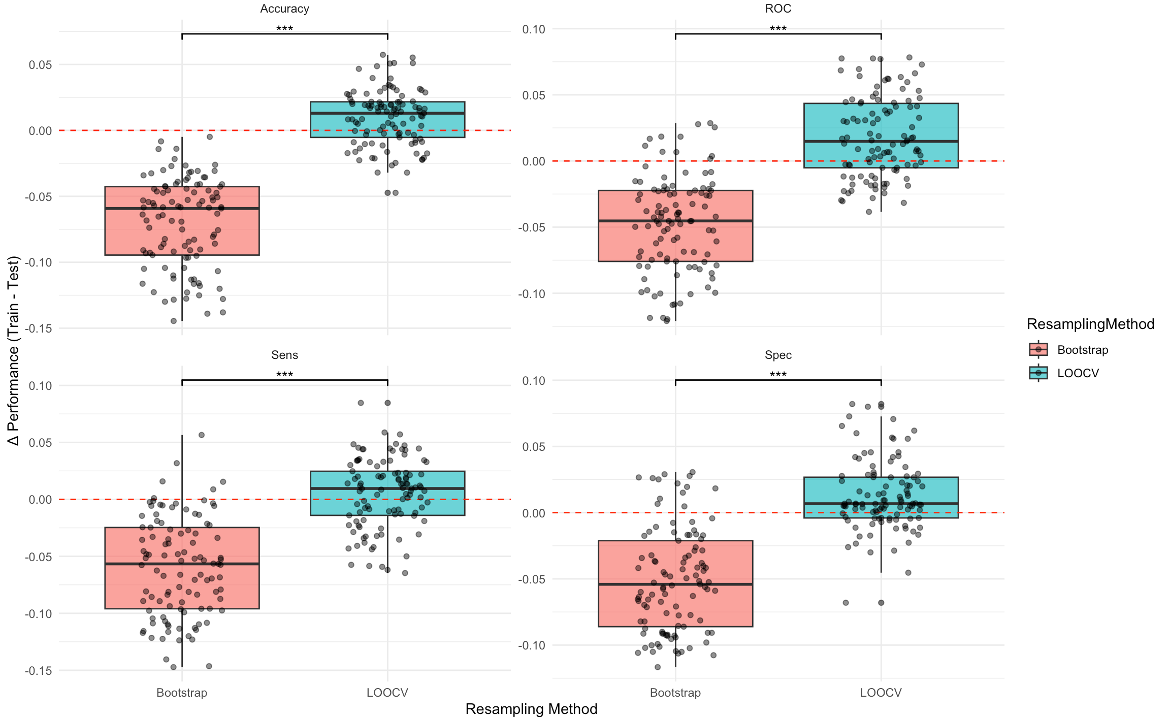

Figure S8: Generalization gap plots of LOOCV and bootstrap models. ***p < 0.001.

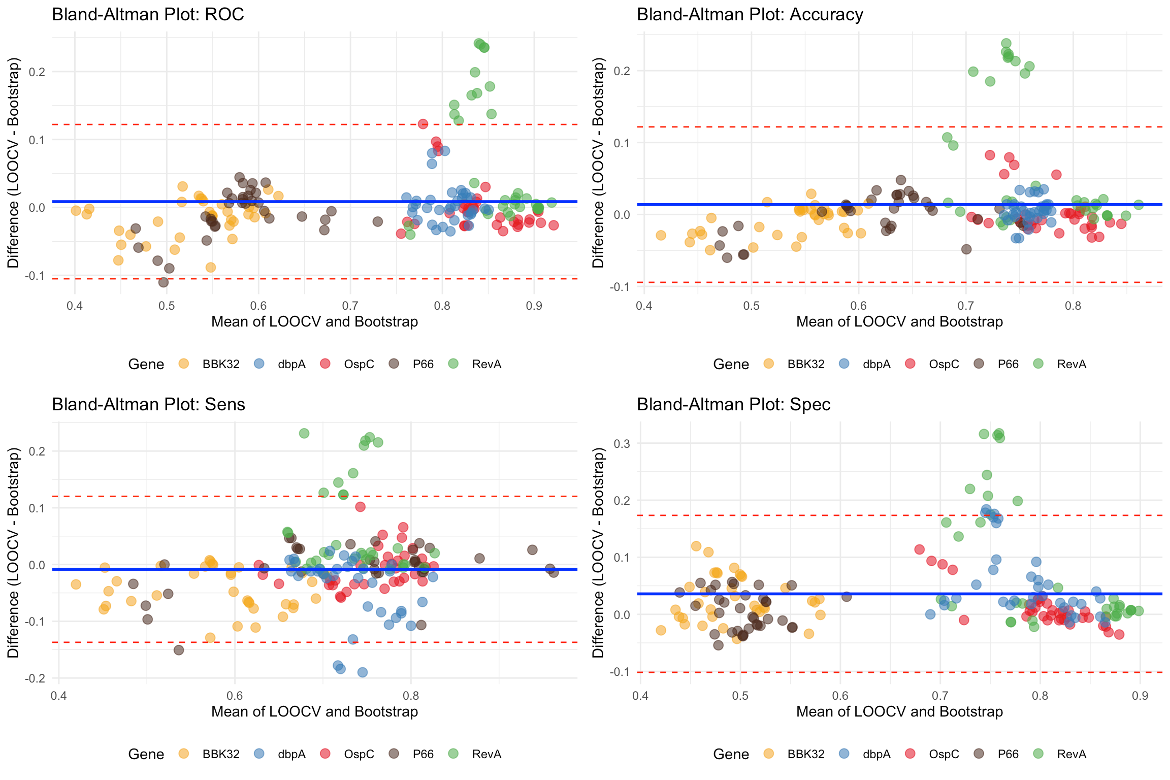

Figure S9: Bland-Altman plots of different performance metrics represent agreements between LOOCV and bootstrap resampling methods.

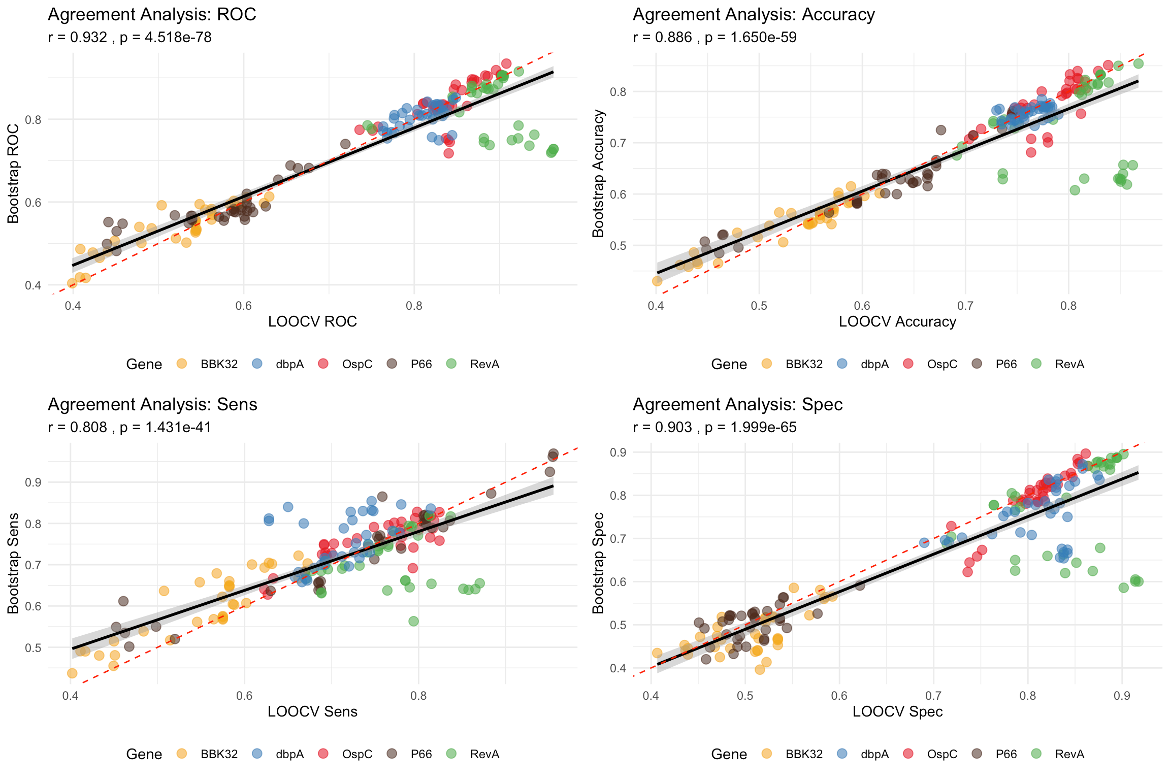

Figure S10: Agreement analysis in different performance metrics between LOOCV and bootstrap models

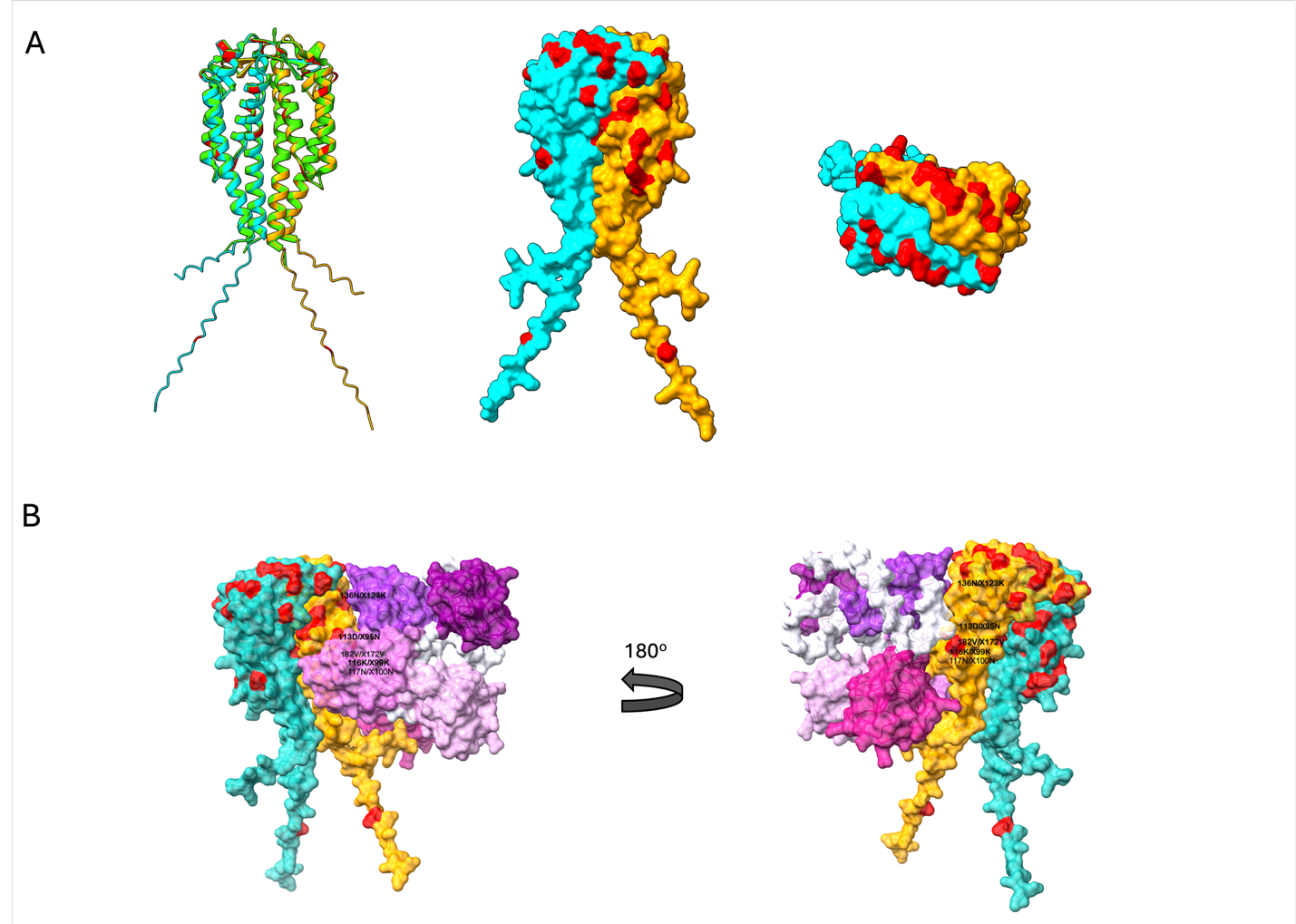

Figure S11: (A) AlphaFold-predicted OspC dimer structures. Left: ribbon representation of the predicted OspC dimer (cyan and yellow), superimposed with the crystal structure of the OspC dimer (green; PDB: 1GGQ). Middle and right: surface views along the 2-fold symmetry axis from different angles. (B). AlphaFold-predicted structure of the plasminogen-OspC-OspC complex. Left and right panels depict the complex in two orientations rotated 180^o^. The five kringle domain of plasminogen are shown in a color gradient from light pink to dark purple. The two OspC subunits are colored cyan and yellow with machine learning-predictive residues highlighted in red.

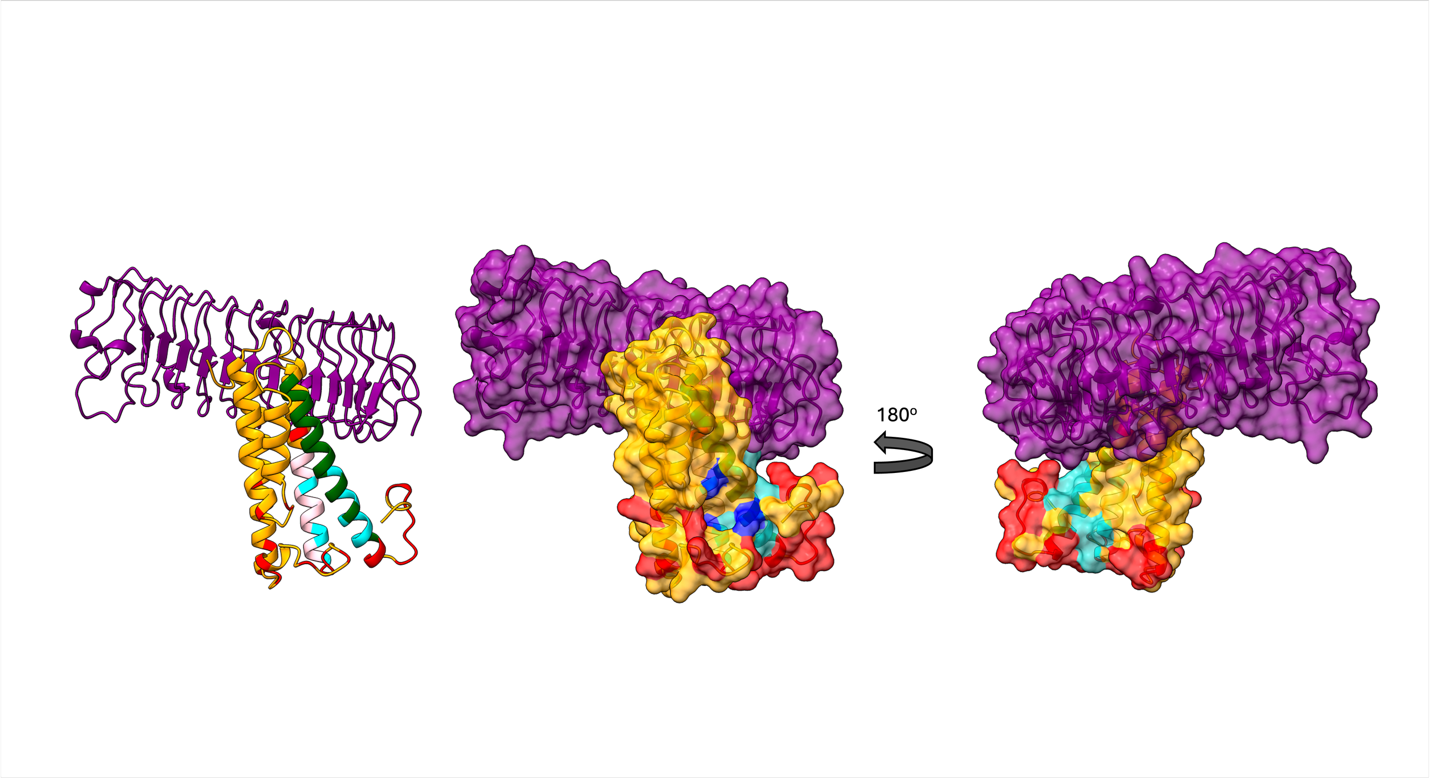

Figure S12: AlphaFold-predicted structure of the decorin-DbpA complex. Left: Ribbon representation of the complex. Middle and right: Surface representations from two orientations (rotated 180^o^). DbpA is shown in yellow and decorin in purple. Residues 76–90 and 152–176 of DbpA, known to be important for decorin binding, are highlighted in pink and green, respectively (95). Red indicates machine learning-identified predictive residues. Cyan marks residues that overlap between ML-predictive sites and known decorin-binding regions. Blue highlights the three conserved Lysines (K82, K163, K170) implicated in glycosaminoglycan binding.

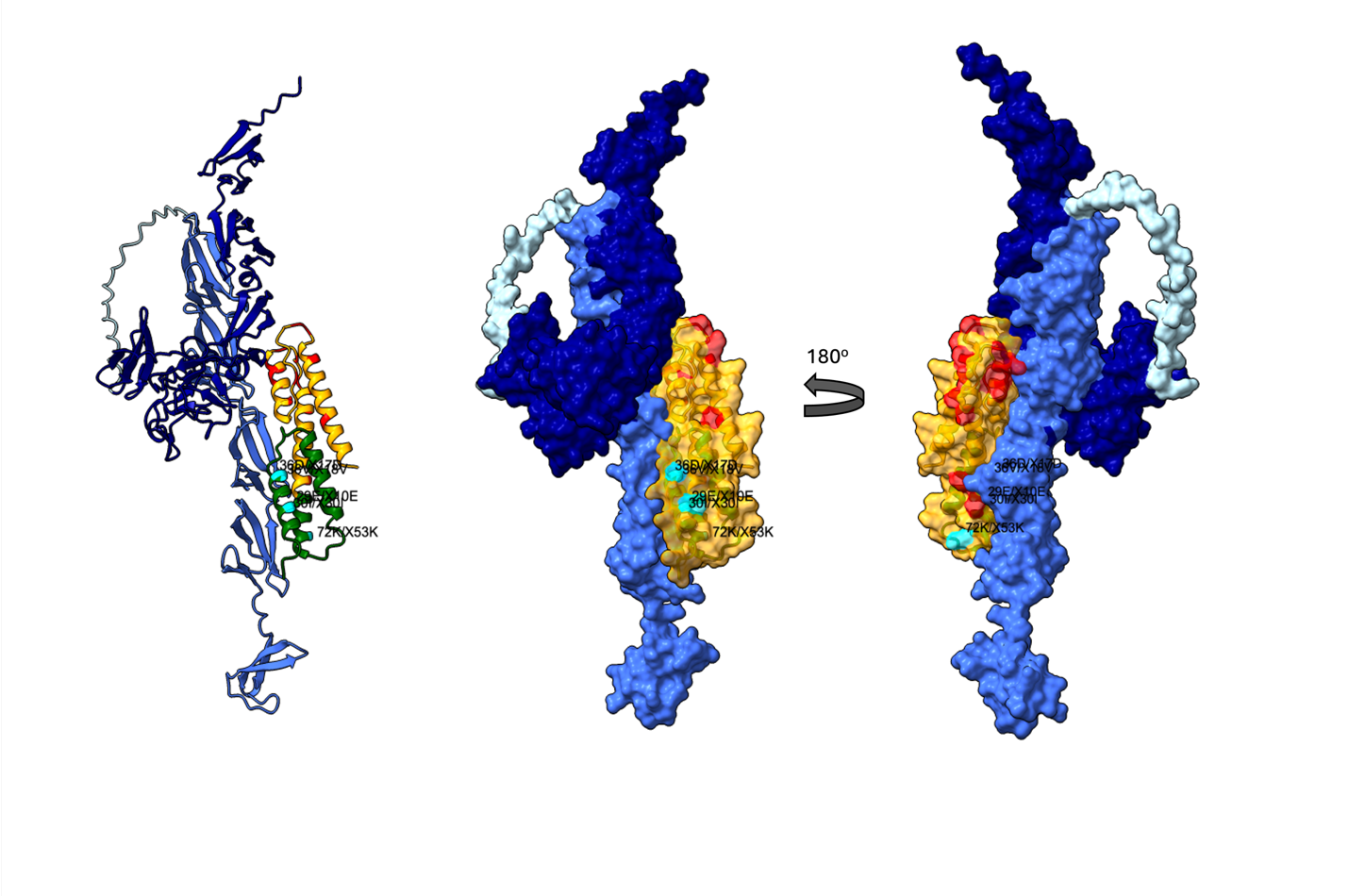

Figure S13: AlphaFold-predicted structure of the fibronectin-RevA complex. Left: Ribbon representation. Middle and right: Surface representations from two orientations (rotated 180^o^). RevA is shown in yellow. Blue and dark blue represent the 30 and 40 kDa regions of fibronectin. Green highlights the N-terminal residues crucial in fibronectin binding (24). Red indicates machine learning-identified predictive residues. Cyan marks ML-predictive residues within the fibronectin-binding N-terminus.

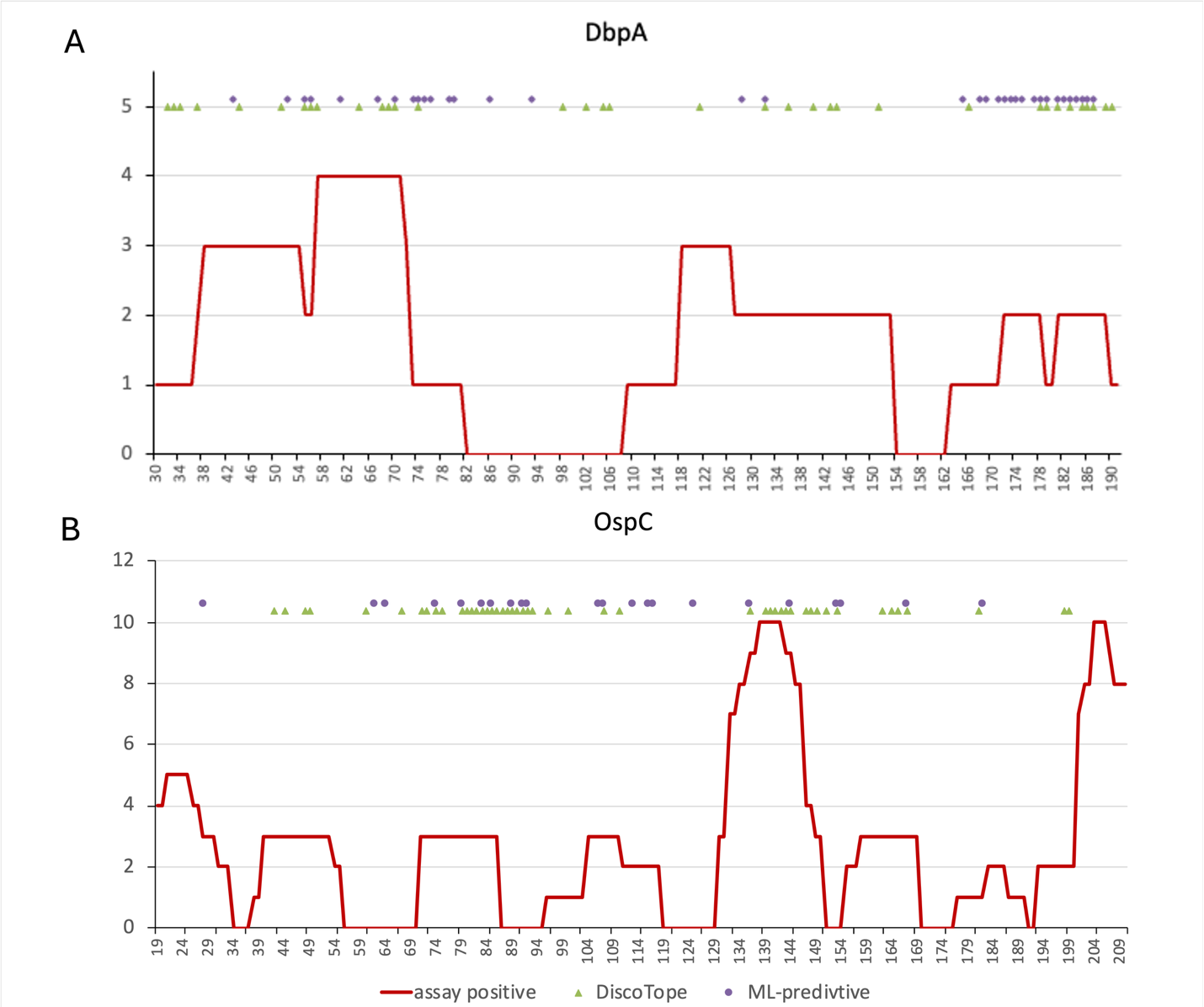
Figure S14: Positive immune assay counts mapped to DbpA and OspC sequence. Data collected from IEDB ImmunoBrowser. Red line indicates the number of positive immune assays mapped to each amino acid position along the protein sequences. Green triangles represent DiscoTope-predicted discontinuous B-cell epitopes, and purple circles indicate machine learning-predicted residues.
